# Circulating Metabolites in Progression to Islet Autoimmunity and Type 1 Diabetes

**DOI:** 10.1101/496943

**Authors:** Santosh Lamichhane, Esko Kemppainen, Kajetan Trošt, Heli Siljander, Heikki Hyöty, Jorma Ilonen, Jorma Toppari, Riitta Veijola, Tuulia Hyötyläinen, Mikael Knip, Matej Orešič

## Abstract

Previous studies suggest that metabolic dysregulation precedes the onset of type 1 diabetes (T1D). However, these metabolic disturbances and their specific role in disease initiation remain poorly understood. Here we analysed polar metabolites from 415 longitudinal plasma samples in a prospective cohort of children in three study groups: those who progressed to T1D (PT1D), who seroconverted to one islet autoantibody (Ab) but not to T1D (P1Ab), and Ab-negative controls (CTR). In early infancy, PT1D associated with downregulated amino acids, sugar derivatives and fatty acids, including catabolites of microbial origin, as compared to CTR. Methionine remained persistently upregulated in PT1D as compared to CTR and P1Ab. Appearance of islet autoantibodies associated with decreased glutamic and aspartic acids. Our findings suggest that children who progress to T1D have a unique metabolic profile, which is however altered with the onset of islet autoantibodies. Our findings may assist in early prediction of T1D.

## Introduction

Type 1 diabetes (T1D) is an autoimmune disease, which arises due to the destruction of the insulin producing pancreatic β-cells by the immune system^1^. The incidence of T1D is highest in children and adolescents in the developed countries^2^ and an increase in disease rate is expected in young children aged less than 5 years^3^. To reverse the increasing rate, early prediction and prevention of T1D is essential. However, the aetiology of T1D disease is complex, multifactorial, and the primary cause for initiation and disease progression is poorly understood^1^. Therefore, predictive and preventive measures for T1D remain unmet medical needs.

Human leukocyte antigen (HLA) complex alleles constitute the most relevant and the strongest genetic risk factor for T1D susceptibility^4^. However, only 3-10% of the individuals with risk HLA loci develop T1D^5^, indicating that exogenous factors such as environmental exposure, diet and gut microbiota likely play a vital role in disease progression^6^. Initiation of β-cell autoimmunity is the first detectable sign of progression towards T1D. However, seroconversion to islet autoantibody positivity may not lead to overt diabetes^7^ and the period between the seroconversion and the appearance of clinical symptoms of T1D may vary between individuals from a few months to many years^8, 9^.

Previous studies suggest that children who progress to T1D have dysregulated metabolic profiles already in infancy^10, 11, 12, 13^, prior to the seroconversion for islet autoantibodies. However, the studies in humans have so far mainly focused on lipids, and there is relatively little information on polar metabolites, such as those involved in central metabolic pathways, in relation to T1D pathogenesis. Herein we study circulating polar metabolite profiles in progression to T1D in a longitudinal study setting.

## Results

### Impact of age on circulating metabolome

We performed metabolomics analysis of polar metabolites in plasma from 120 children, divided into three study groups: those who progressed to T1D (PT1D, n = 40), who seroconverted to at least one autoantibody (Ab) positivity but without clinical symptoms of T1D (P1Ab, n = 40), and matched Ab negative controls (CTR, n = 40). For each participant, plasma samples were collected corresponding to the ages of 3, 6, 12, 18, 24, and 36 months (**Fig. 1**). We detected metabolites from across a wide range of chemical classes including amino acids, carboxylic acids (mainly free fatty acids and other organic acids), hydroxyacids, phenolic compounds, alcohols, and sugar derivatives.

Principal components analysis (PCA)^14^ of the complete dataset including identified metabolites displayed an age-dependent pattern **(Supplementary information (SI) Fig.1**). To resolve the impact of age on plasma metabolome, we performed analysis of variance (ANOVA)-simultaneous component analysis (ASCA) ^15^ by incorporating three factors: age, gender, study cases (CTR, P1Ab, PT1D) and their interactions. As expected, age related variation displayed maximum effect (4.2 %, p= 0.001) in the circulating metabolome as compared to the impact of the other two factors, ‘study groups’ (1.2 %, p = 0.001) and ‘gender’ (0.5 %, p = 0.002). Notably, the interaction factor ‘age and cases’ also showed a significant effect (2.9 %, p = 0.033), while interactions between other factors (age/gender and case/gender) remained insignificant (p = 0.508 and p = 0.221, respectively).

**Figure 1.**
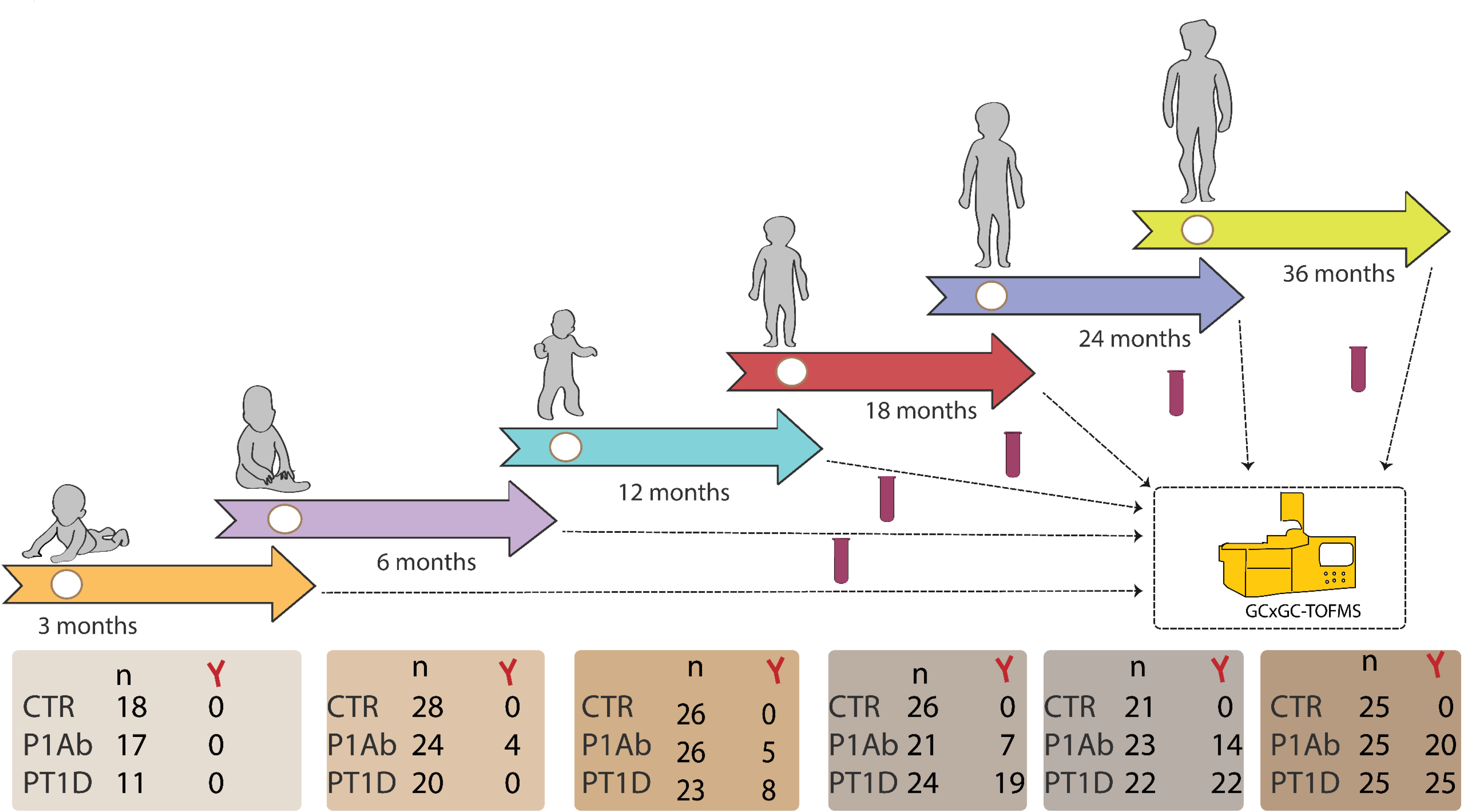
An overview of the study design. The study cohort comprises the samples from children, who progressed to T1D (PT1D), who seroconverted to one islet autoantibody but did not progress to T1D during the follow-up (P1Ab), and control (CTR) subjects who remained islet autoantibody negative during the follow-up until the age of 15 years. For each child, longitudinal plasma samples were drawn, corresponding to the ages of 3, 6, 12, 18, 24, and 36 months. In each age cohort and study group, number of autoantibody positive children is marked and represented with Y-shape.

The scores from the first principle component (PC1) of the factor ‘age’ clearly showed an age-related trajectory in the circulatory metabolites (**Fig. 2**). The loading revealed high levels of branched chain amino acids (BCAA) in the 18, 24 and 36 month age-cohorts, whereas tryptophan, 3-indole acetic acid (tryptophan derivative) and carboxylic acids (mainly free fatty acids) were elevated during early infancy (3 and 6 months). However, we did not detect any age-dependent patterns in phenolic compounds, alcohols, hydroxyacids, and sugar derivatives (**SI Fig. 2**).

**Figure 2.**
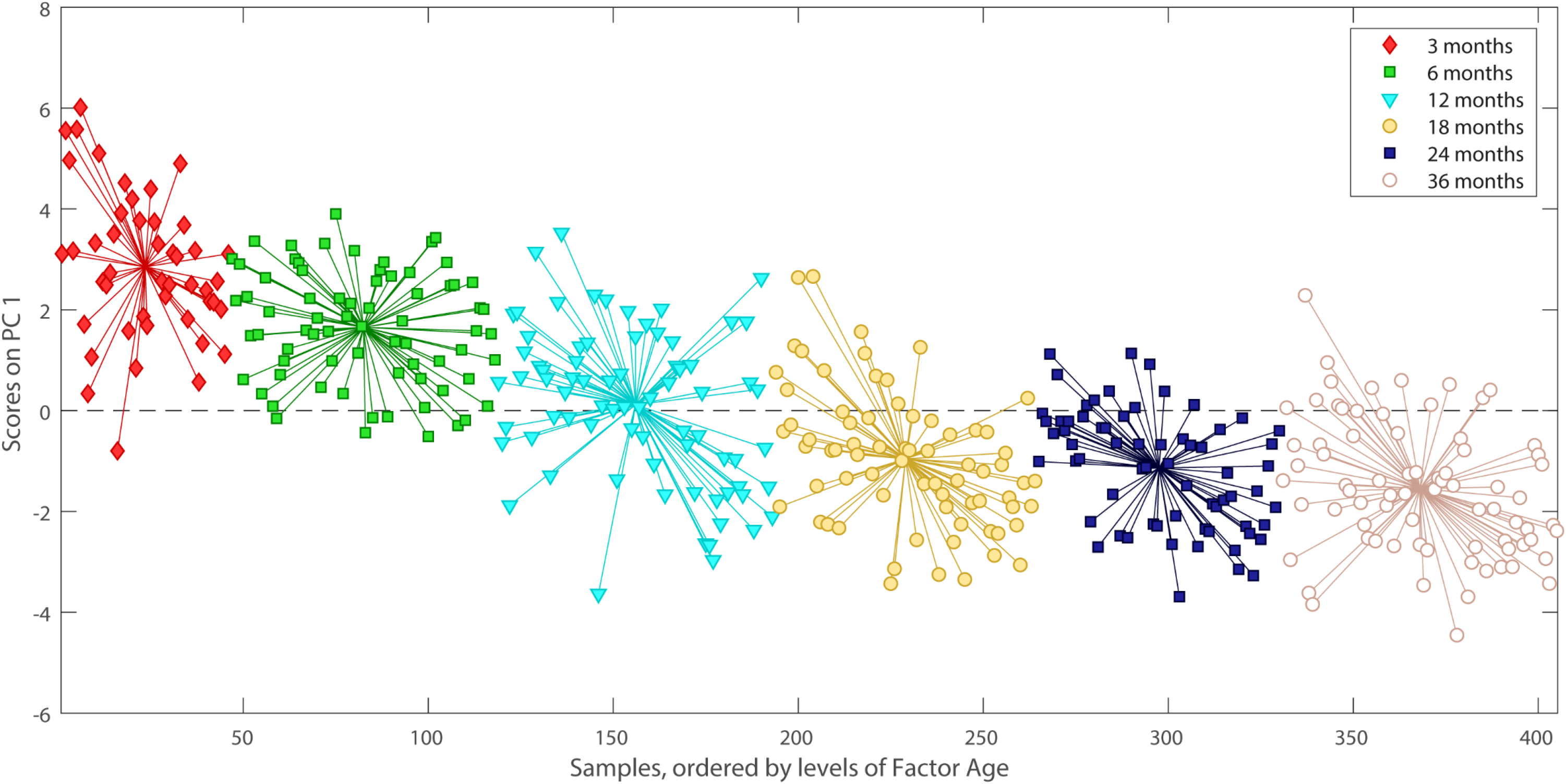
PCA score plots of the factor age, based on ASCA. These scores represent the metabolomics dataset arranged according to the age in the PCA score plot. Each sample is represented by a point and coloured according to the age. The ages of the participants are marked on the x-axis while y-axis represents the sample score. Samples with similar score cluster together.

### Metabolite profiles during progression to islet autoimmunity and T1D

Considering the age as a major confounder in the plasma metabolome, we performed age-matched comparisons between the three study groups (CTR, P1Ab, and PT1D). Univariate analysis revealed that all major metabolite classes, including amino acids, free fatty acids and sugar derivatives were altered, already in infancy, among the children who later progressed to T1D (**Fig. 3**). Altogether 15 metabolites were different between PT1D and CTR groups at three months of age (nominal p-value < 0.05). Nine out of 15 metabolites were significantly lower in T1D progressors as compared to controls (FDR threshold of 0.1) (Fig. 3, **SI Table 1**). In order to assess if gender had an impact on plasma metabolite levels in children at three months of age, we carried out ASCA analysis with factor: study cases and gender, and their interaction. When evaluating the statistics from these factors, we found only study cases had significant effect (p = 0.012), while gender and their interaction remained insignificant (p = 0.081 and p = 0.73, respectively). The score of the factor ‘study cases’ showed distinct metabolic clusters between PT1D, P1Ab and CTR, suggesting that specific metabolic changes precede islet autoimmunity and T1D. The loadings disclosed that methionine, 2-ketoisocaproic acid, bisphenol A, pyruvic acid, glycerol-2-phosphate, and levoglucosan were higher in the PT1D group when compared with the P1Ab and CTR groups (**SI Fig. 3**).

**Figure 3.**
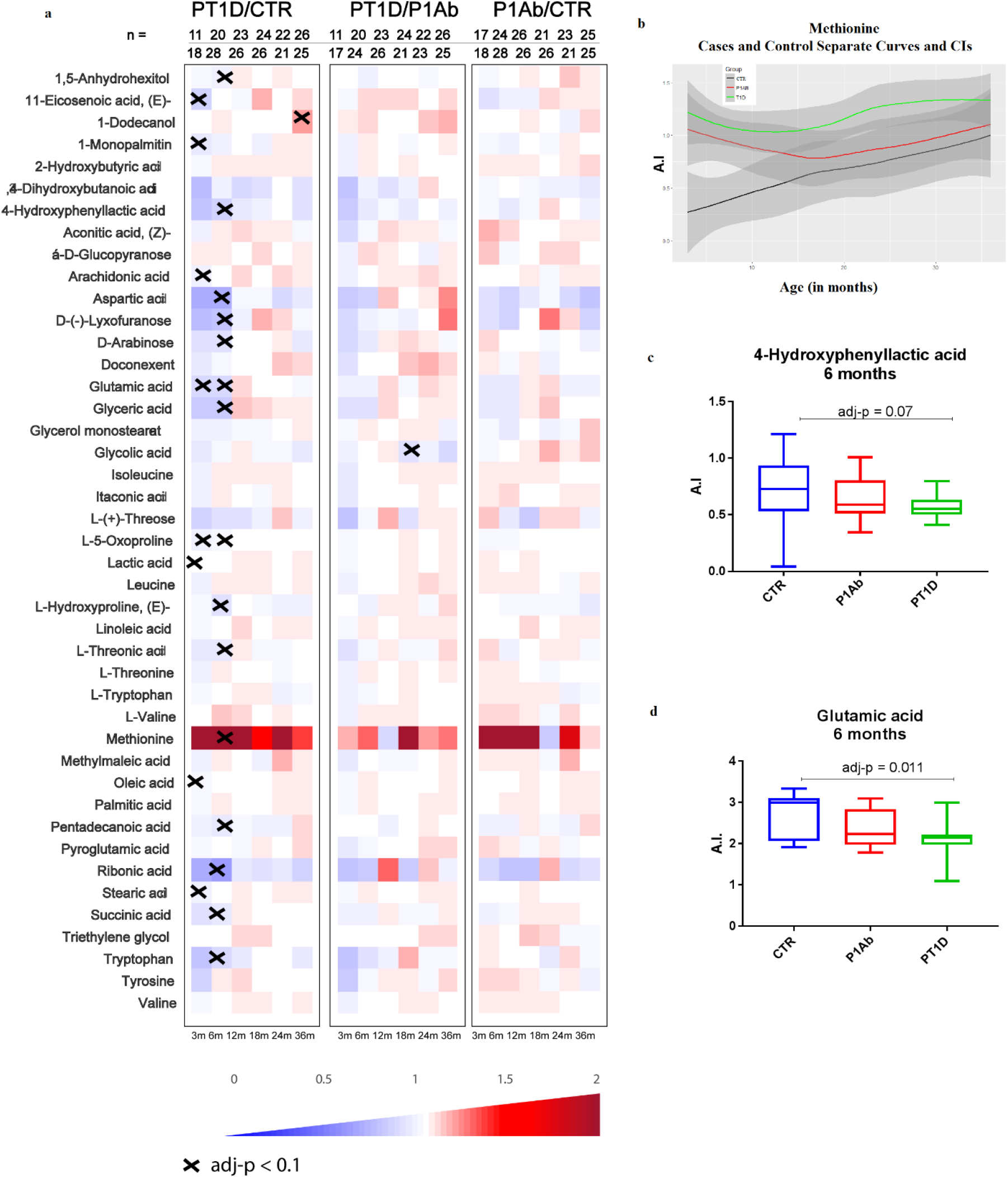
Comparison of metabolomes in three study groups in different age cohorts. **(a)** Heat map showing 43 metabolites representative of different metabolic classes that change between PT1D, P1Ab and CTR. Differences in metabolite concentrations were calculated by dividing mean concentration in PT1D by the mean concentrations in P1Ab and CTR. **(b)** The *loess* curve plot of methionine concentration in time for the three study groups. **(c)** Concentration of 4-hydroxyphenyllactic acid at 6 months of age. **(d)** Concentration of Glutamic acid at 6 months of age. **X** represents the adjusted p-values < 0.1.

At 6 months of age, altogether 20 metabolites differed between PT1D and CTR (nominal p-value < 0.05). Fifteen of these circulating metabolites passed the FDR threshold of 0.1 (**Fig. 3a-c**, **SI Table 2**), including several amino acids, sugar derivatives, free fatty acids and various other organic acids. The levels of most of these metabolites decreased in T1D progressors during the same period as compared to CTR. Only methionine was found increased in PT1D as compared to CTR at the age of 6 months. In addition, multivariate ASCA analysis revealed that only study group (CTR, P1Ab, and PT1D) had a significant effect (p = 0.004) in the plasma metabolites of 6-month-old children, whereas the impact of gender (p =0.180) and its interaction with study group (p = 0.269) remained insignificant.

Next, we sought to examine weather children across the three study groups had altered plasma metabolite levels in the age cohorts of 12, 18, 24, and 36 months. With the exceptions of 1-dodecanol and glycolic acid, no other statistically significant differences between the study groups were observed (FDR threshold of 0.1). At 36 months of age, dodecanol level was higher in PT1D as compared to CTR. Meanwhile, glycolic acid was lower in PT1D as in P1Ab at 18 months of age. However, in longitudinal series these metabolites showed inconsistent trends (**Fig. 3b**).

We also studied whether group of metabolites at early age associated with a specific metabolic pathway. The altered metabolites (p < 0.05) between CTR and PT1D at 3 and 6 months of age were subjected to metabolic pathway analysis (MetPA) in MetaboAnalyst^16^. In line with findings at the individual metabolite levels, we found that four metabolic pathways, linoleic acid metabolism, arachidonic acid metabolism, alanine, aspartate and glutamate metabolism and D-glutamine and D-glutamate metabolism remained altered between PT1D and CTR groups at the age of three months (**Fig. 4a**, **SI Table 3**). Similarly, at 6 months of age, MetPA revealed that alanine, aspartate and glutamate metabolism, D-glutamine and D-glutamate metabolism, tryptophan metabolism, arginine and proline metabolism, as well as aminoacyl-tRNA biosynthesis remained dysregulated between the controls and T1D progressors (**Fig. 4b**, **SI Table 4**).

**Figure 4.**
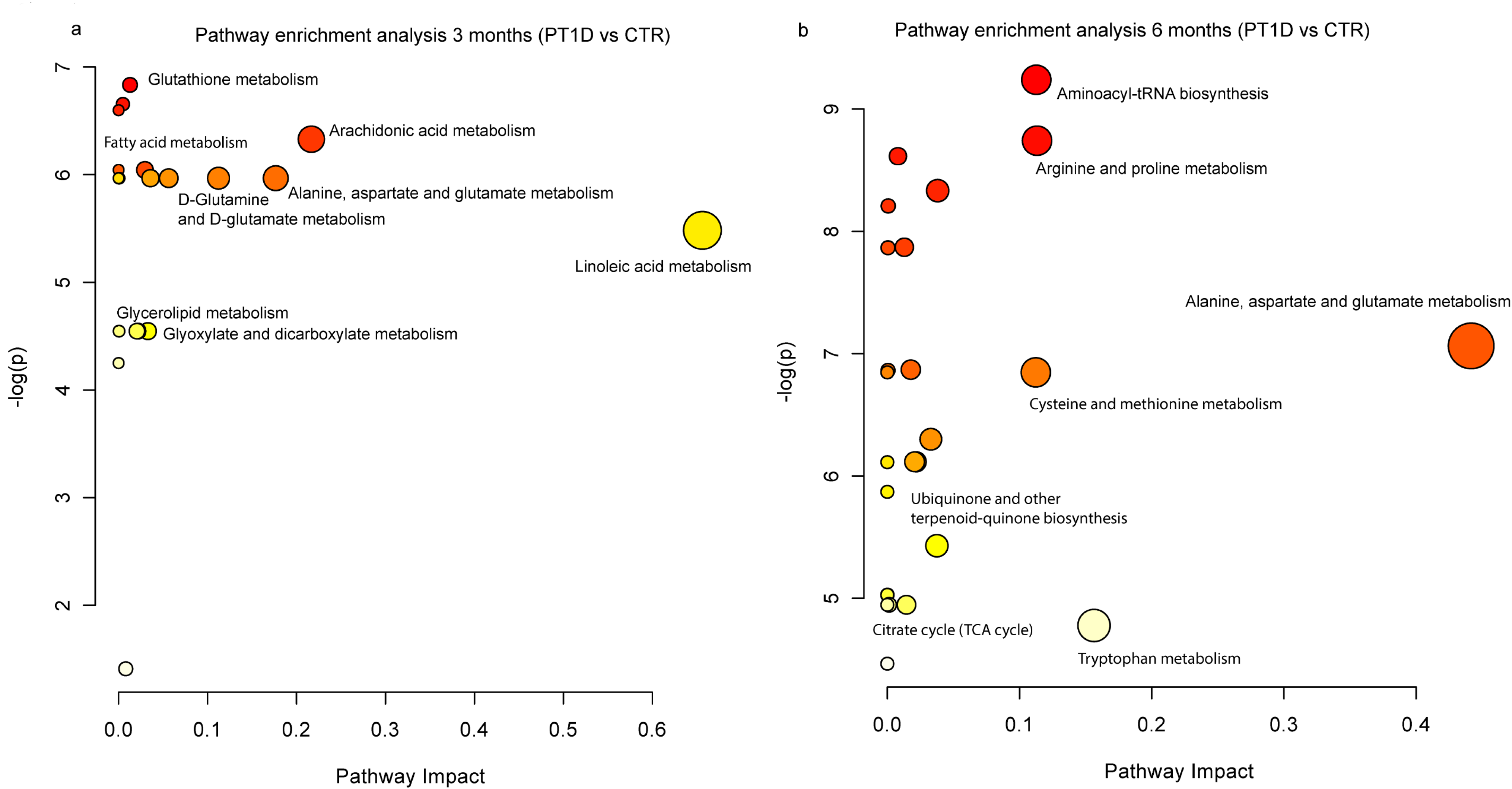
Pathway analysis of significantly different metabolites between CTR and PT1D at **(a)** 3 and **(b)** 6 months of age. The pathways are shown according to the p values from the pathway enrichment analysis and pathway impact values from the pathway topology analysis. The metabolic pathways with impact value > 0.1 were considered the most relevant pathways involved. Pathway impact values were calculated from pathway topology analysis using MetaboAnalyst.

**Figure 5.**
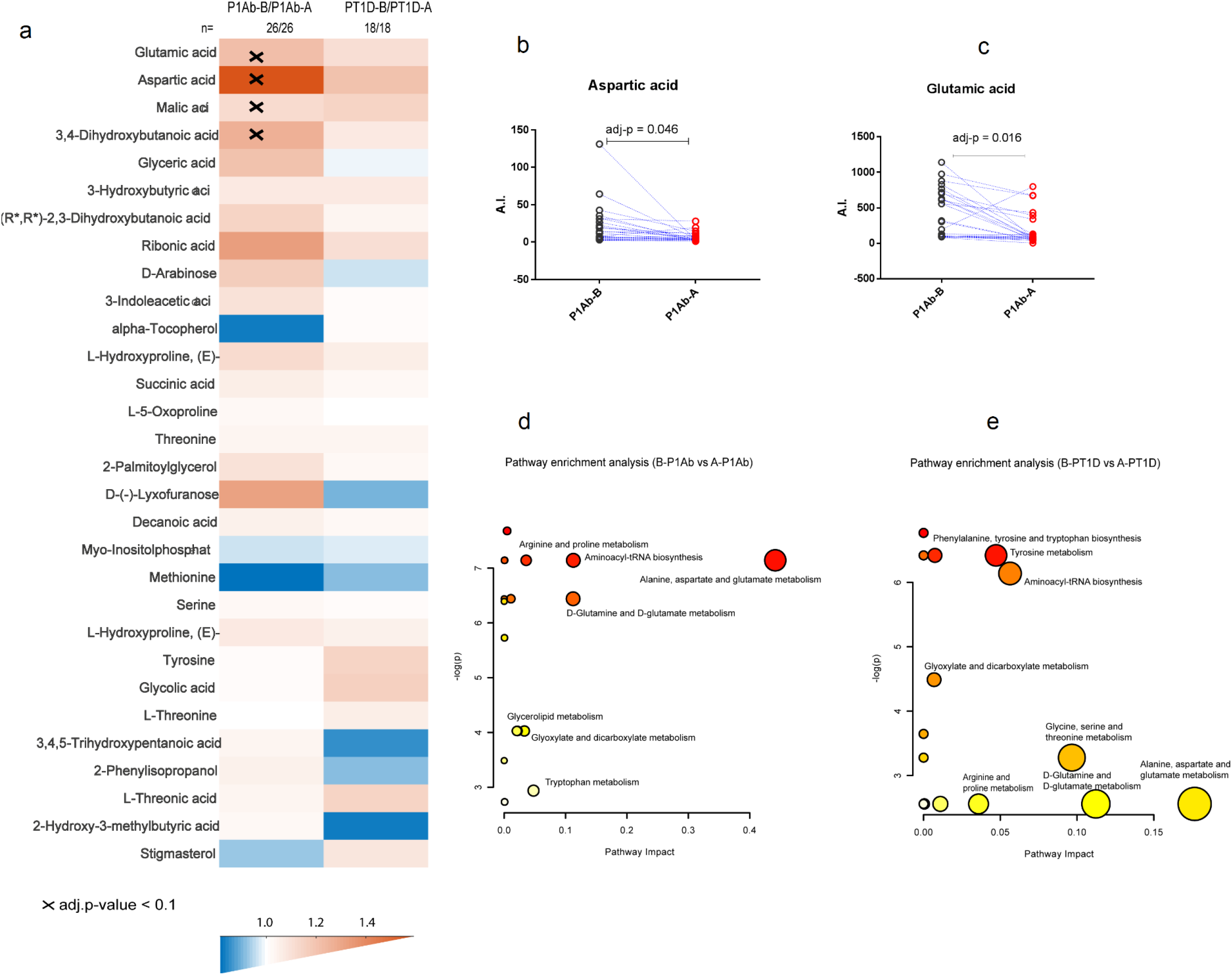
The effect of islet autoantibody positivity on metabolite profiles. **(a)** The most discriminating metabolites between the last available samples obtained before the first islet autoantibody appeared and the first available samples after the emergence of the first islet autoantibody, in P1Ab and PT1D groups. The pairwise scatter plot of **(b)** aspartic and **(c)** glutamic acid before and after the first appearance of islet autoantibodies. Pathway analysis of differentially expressed metabolites between **(d)** B-P1Ab & A-P1Ab, and **(e)** B-PT1D & A-PT1D. Top pathways identified include Alanine, aspartate and glutamate metabolism. Abbreviations: Before seroconversion in P1Ab (B-P1Ab), after seroconversion in P1Ab (A-P1Ab), before seroconversion in progressors (B-PT1D), after seroconversion in progressors (A-PT1D).

### Metabolome before and after the first appearance of islet autoantibodies

In order to study the effect of islet seroconversion on metabolome, we compared metabolite levels before and after the appearance of first islet autoantibody in P1Ab and PT1D groups. Pairwise comparison revealed that eleven metabolites were altered by seroconversion in P1Ab (nominal p-value < 0.05, **SI Table 5**), with four passing the FDR threshold of 0.1 (glutamic, aspartic, malic, and 3, 4-dihydroxybutanoic acids) (**Fig. 4**). We detected seven metabolites altered before and after islet autoantibody appearance in PT1D (nominal p-value < 0.05), but none of these passed the FDR threshold of 0.1 (**SI Table 6**). Metabolic pathway analysis corroborated these findings and revealed that alanine, aspartate and glutamate metabolism were altered when comparing the pathways before and after seroconversion within P1Ab and PT1D groups (**Fig.4**). However, the level of impact for these pathways varied between P1Ab and PT1D, with impact values 0.441 and 0.176, respectively. Other relevant pathways and their impact are summarized in **SI Table 7** and **Table 8**. When examining metabolite level changes in relation to the appearance of specific islet autoantibodies (islet cell antibodies (ICA), insulin autoantibodies (IAA), islet antigen 2 autoantibodies (IA-2A), and GAD autoantibodies (GADA)), no specific associations were identified, which may be due to the small number of cases per individual autoantibody.

## Discussion

Our study identified specific metabolic disturbances in children who progressed to T1D, as compared to their age matched controls including children who developed a single islet autoantibody but did not progress to T1D during the follow-up. We found that such metabolic dysregulation exists before the first signs of islet autoimmunity. In agreement with earlier studies^10, 17, 18^, a strong association of the metabolome was observed with age. We identified a distinct plasma amino acid profile in PT1D children, particularly at the ages of 3 and 6 months. Glutamic and aspartic acids as well as tryptophan remained downregulated during the early infancy in PT1D as compared to CTR, but not to P1Ab. In our previous study of polar metabolites in T1D progression, we found no significant difference in different age cohorts when comparing PT1D and CTR groups^13^, which may however be attributable to the small number of individuals in the metabolomics part of that study. Notably, in agreement with the previous study, we also observed that the appearance of islet cell autoantibodies was associated with down-regulation of aspartic and glutamic acids^13^, also corroborated by observed change in alanine, aspartate and glutamate metabolism in the MetPA. Our findings are consistent with previous study suggesting that amino acid dysregulation precedes the appearance of islet autoantibodies and progression to T1D ^12^. Several free fatty acids were also downregulated at 3 months of age. During basal metabolic processes, triglycerides (TGs) are broken down to fatty acid and glycerol^19^. Fatty acid act as an important fuel source for cells, which is required to maintain systematic energy homeostasis^20^. Usually, under conditions when the availability of carbohydrate is not enough, the fatty acids are alternative substrate for energy production^21^. Here, fatty acid decrease may be an indication of increased energy demand in PT1D, further substantiated by the diminishment of circulating sugar derivatives as well as altered linoleic acid metabolism and arachidonic acid metabolism. This is also in line with our previous report^10^ associating downregulated TGs and phospholipids in the PT1D group, supporting the view that altered energy metabolism is involved in the initiation of the autoimmune process and T1D.

Accumulating evidence suggests that perturbations in the gut microbial structure are associated with, and contribute to the pathogenesis of β-cell autoimmunity and to overt T1D^22,23, 24^. Here we found that 4-hydroxyphenyllactic acid^25, 26^, 11-eicosenoic acid^27^, and succinic acid^28^, the metabolites of potential microbial origin (catabolites), are significantly downregulated at early age (3 and 6 months) in PT1D. The tryptophan derived microbial catabolite 3-indoleacetic tended to be also downregulated in PT1D (**SI Fig. 4**). Catabolites generated by the gut microbes are vital to the intestinal homeostasis^26, 29^, thus it is likely that scarcity of substrates for microbial catabolism contribute to the dysbiosis associated with progression to T1D.

While most of the amino acids were downregulated in PT1D as compared to CTR and P1Ab, methionine remained persistently upregulated in T1D progressors. This appears to be in disagreement with previous studies in BABYDIAB and MIDIA cohorts, which showed decreased level of methionine in autoantibody positive individuals and T1D progressors, respectively^18, 30^. This discrepancy may however be explained: (1) BABYDIAB study compared children seroconverting early in life (≤2 years) to those who developed autoantibodies at older age, while (2) MIDIA study highlighted differences, which were mainly linked to the age of the children and the duration of breastfeeding^30^. We performed similar comparison to that of BABYDIAB in the current study setting but found no significant differences between the groups compared. The observed differences suggest disrupted methionine metabolism in PT1D. Methionine can be salvaged endogenously by protein/homocysteine degradation, polyamine synthesis, or by transsulfuration pathway^31^, and the disturbances in these pathways could modulate the neonatal epigenetic processes including the DNA methylation and chromatin remodelling and consequently influence various immunologic responses^32^.

The ASCA multivariate analysis revealed that plasma BPA was upregulated in PT1D group, although univariate analysis across different age cohorts did not reveal significant changes between the groups. Studies in experimental model of autoimmune diabetes suggest that increased BPA exposure is associated with accelerated development of autoimmune diabetes^33, 34^. However, we consider that at the present stage our findings on the association of BPA and T1D are still inconclusive, because (1) in our study setting we could not control for the effect of sample storage on the plasma BPA levels and (2) the levels of BPA were not quantified. Clearly further studies in clinical settings are merited in order to establish the effect of exposure to BPA and other environmental toxicants on the progression of T1D or other autoimmune diseases.

Taken together, while confirming several earlier findings, the present study highlights the importance of core metabolic pathways such as amino and fatty acid metabolism in early pathogenesis of T1D. Metabolites of microbial origin were also found associated with T1D progression. We also observed that appearance of islet autoantibodies does have an effect on the amino acid levels, specifically on glutamic and aspartic acids. However, these changes do not seem to be specifically associated with T1D but are instead a general feature of islet autoimmunity - suggesting that amino acid imbalance may be a contributing factor in the initiation of autoimmunity^13^. Our study also indicates that the largest metabolic changes associated with T1D progression occur already in early infancy, then these early metabolic signatures become less pronounced or even disappear with age, particularly after the initiation of islet autoimmunity. This may have important implications in the search of early metabolic markers of T1D and for understanding the disease pathogenesis.

## Methods

These methods are expanded versions of descriptions in our related work^10^.

### Study setting

The plasma samples were from the Finnish Type 1 Diabetes Prevention and Prediction Study (DIPP) ^35^. The DIPP study has screened more than 220,000 newborn infants for HLA-conferred susceptibility to T1D in three university hospitals (Turku, Tampere, and Oulu) in Finland ^36^. The subjects in the current study were from the subset of DIPP children from the Tampere study centre. The ethics and research committee of the participating university hospital approved the study protocol and the study fallowed the guidelines of the Declaration of Helsinki. Parent for all participants signed written informed consent at the beginning of the study. We collected five longitudinal samples per child, corresponding to either of the ages of 3, 6, 12, 18, 24, and 36. This longitudinal cohort comprises of samples from 120 children: 40 progressors to T1D (PT1D), 40 who tested positive for at least one Ab in a minimum of two consecutive samples but did not progress to clinical T1D during the follow-up (P1Ab), and 40 controls (CTR) subjects who remained islet autoantibody negative during the follow-up until the age of 15. We matched the participants in the three study group for HLA-associated diabetes risk, gender and period of birth. In total, we collected 415 non-fasting, blood samples. We separated plasma within 30 minutes after the blood collection by centrifugation at 1600g for 20 minutes at room temperature. The plasma samples were stored at -80°C until analysed.

### HLA genotyping

HLA-conferred susceptibility to T1D was analysed using cord blood samples as described by Nejentsev *et al.* ^37^. Briefly, the HLA-genotyping was performed with time-resolved fluorometry based assay for four alleles using lanthanide chelate labelled sequence specific oligonucleotide probes detecting DQB1*02, DQB1*03:01, DQB1*03:02, and DQB1*06:02/3 alleles^38^. The carriers of the genotype DQB1*02/DQB1*03:02 or DQB1*03:02/x genotypes (here x≠ DQB1*02, DQB1*03:01, DQB1*06:02, or DQB1*06:03 alleles) were categorized into the T1D risk group and recruited for the DIPP follow-up program.

### Detection of islet autoantibodies

The participants with HLA-conferred genetic susceptibility were prospectively observed for the appearance of T1D associated autoantibodies (islet cell antibodies (ICA), insulin autoantibodies (IAA), islet antigen 2 autoantibodies (IA-2A), and GAD autoantibodies (GADA). These autoantibodies were analysed in the Diabetes Research Laboratory, University of Oulu from the plasma samples taken at each follow-up visit as described ^39^. ICA antibodies were detected with the use of indirect immunofluorescence, whereas the other three autoantibodies were quantified with the use of specific radiobinding assays^40^. We used cut-off limits for positivity of 2.5 Juvenile Diabetes Foundation (JDF) units for ICA, 3.48 relative units (RU) for IAA, 5.36 RU for GADA, and 0.43 RU for IA-2A. The disease sensitivity and specificity of the assay for ICA were 100% and 98%, respectively, in the fourth round of the international workshops on standardization of the ICA assay. According to the Diabetes Autoantibody Standardization Program (DASP) and Islet Autoantibody Standardization Program (IASP) workshop results in 2010-2015, disease sensitivities for the IAA, GADA and IA-2A radio binding assays were 36-62%, 64-88% and 62-72%, respectively. The corresponding disease specificities were 94-98%, 94-99% and 93-100%, respectively.

### Analysis of polar metabolites

After randomization and blinding, 415 plasma samples were used for extraction. Plasma was thawed on ice and aliquoted. 30 μl of aliquot was used for analysis of polar metabolites. Extraction was performed with 400 μl of methanol as previously described ^41^. For quality control and normalization purpose 10 μl of following group-specific internal standard mix was added into extraction solvent. Internal standard mix was composed of: heptadecanoic acid-d33 (175.36 mg/l), valine-d8 (35.72 mg/l), succinic acid-d4 (58.54 mg/l) and glutamic acid-d5 (110.43 mg/l). Internal standards were purchased from Sigma-Aldrich (Steinheim, Germany) and methanol from Honeywell Riedel de Haën (Seezle, Germany). Samples were vortexed and left to precipitate for 30 min on ice. After protein precipitation, extracts were centrifuged (Eppendorf; 5427R) for 3 min on 10000 rpm. 180 μl of supernatant was transferred into GC vials and stored for further use. Same procedure was applied for clinic-pooled plasma which was used for quality control and batch correction. Calibration curves were made from the following standards: Fumaric acid, Aspartic acid, Succinic acid, Malic acid, Methionine, Tyrosine, Glutamic acid, Phenylalanine, Arachidonic acid, Isoleucine, 3-Hydroxybutyric acid, Glycine, Threonine, Leucine, Proline, Serine, Valine, Alanine, Stearic acid, Linoleic acid, Palmitic acid and Oleic acid. Standards were purchased from Sigma-Aldrich (Steinheim, Germany) and dissolved in methanol. Calibration curves included at least six concentration points in the range from 1 ng/sample up to 3000 ng/sample, depending on the abundance in plasma. R^2^ was from 97.1% up to 99.9%.

Derivatization was performed instrumentally using MPS2 (Gerstel; Mülheim an der Ruhr, Germany) with two robotic hands guided by Maestro software. Samples were evaporated to dryness before two-step extractions. In the first step 25 μl of methoxyamine hydrochloride (TS-45950; Thermo Scientific: USA) was added to the sample. While mixing, the solution was incubated for one hour at 45 °C. In the second step, 25μl of N-methyl-N-trimethylsilyltrifluoroacetamide (Sigma-Aldrich; Steinheim, Germany) was added. Incubation was again performed for one hour at 45 °C. Before injection 50 μl of hexane was added to increase the volatility of the solvent. Additional standards here added during derivatization. n-alkanes (c = 8 mg/l in MSTFA) were used for calculation of retention indexes and 4,4’-dibromooctafluorobiphenyl (c = 9.8 mg/l in hexane) were used as syringe standard to control the quality of injection. 1 μl of derivatized sample was injected after derivatization program was completed.

Derivatised compounds were analysed using Pegasus 4D system (LECO; Saint Joseph; USA). Method used is based on two-dimensions gas chromatography followed by high speed time of flight acquisition of EI fragmented mass spectra. Primary column was 10 m × 0.18 mm I.D. Rxi-5 ms (Restek Corp., Bellefonte, PA, USA) and secondary column 1.5 m × 0.1 mm I.D. BPX-50 (SGE Analytical Science, Austin, TX, USA). System was guarded by retention gap column from deactivated silica (1.7m, 0.53 mm ID, FS deactivated, Agilent technologies, USA). Modulator used nitrogen gas which was cryogenically cooled. Second dimension cycle was 4s. Temperature program started with 50 °C (2 min) then a gradient of 7°C up to 240°C was applied and finally 25°/min to 300 °C where it was held stable for 3 min. Temperature program of secondary column was maintained 20 °C higher than the primary column. Acquisition rate was kept on 100 Hz. Instrument was guided by ChromaTOF software (version 4.32; LECO Corporation, St. Joseph, USA) which was also used calculating area under the peaks with SN>100 and potential identification of peaks using NIST14 and in-house library. Processing method included calculation of retention indexes. Selected compounds were quantified against external calibration curves. Results were exported as text files for further processing with Guineu^42^ software.

### Data analysis

All statistical analyses were performed on log-transformed intensity data. The transformed data were mean cantered and auto scaled prior to multivariate analysis. The multivariate analysis was done using the PLS Toolbox 8.2.1 (Eigenvector Research Inc., Manson, WA, USA) in MATLAB 2017b (Mathworks, Inc., Natick, MA, USA). PCA was initially performed to highlight trend and to get an overview of variation in the dataset. ANOVA-simultaneous component analysis (ASCA) a multivariate extension of ANOVA analysis was performed to allow interpretation of the variation induced by the different factors including age, sex, case, and their interaction^15^.

Wilcoxon rank-sum test was performed for comparing the two study groups of samples (e.g. PT1D vs. P1Ab) in a specific age cohort. For comparison, one sample per subject, closest to the age within the time window, has been used in each test. Paired t-test was performed for the matched groups of samples (e.g. before vs. after seroconversion). The resulting nominal p-values were corrected for multiple comparisons using Benjamin and Hochberg approach^43^. The adjusted p-values < o.1 (q-values) were considered significantly different among the group of hypotheses tested in a specific age cohort. All of the univariate statistical analyses were computed in MATLAB 2017b using the statistical toolbox. The fold difference was calculated by dividing the mean concentration of a lipid species in one group by another, for instance mean concentration in the PT1D by the mean concentration in P1Ab, and then illustrated by heat maps. The locally weighted regression plot was made using smoothing interpolation function loess (with span = 1) available from ggplot2^44^ package in R^45^. The individual lipids levels were visualized as box plot using GraphPad Prism 7 (GraphPad Software, La Jolla, CA, USA).

Pathway analysis of the significant metabolites (nominal p-values < 0.05) was performed using metabolomics pathway analysis (MetPA) tool in MetaboAnalyst 4.0^16^. The compounds unmatched during compound name matching were excluded from the subsequently pathway analysis. We implemented Globaltest hypergeometric testing method for the functional enrichment analysis. The pathway topological analysis was based on the relative betweenness measures of a metabolite in a given metabolic network and for calculating the pathway impact score. Based on the impact values from the pathway topology analysis the impact value threshold was set to > 0.10.

## Data availability

The metabolomics data and the associated meta-data are deposited at the MetaboLights database ^46^ with the acquisition number (MTBLS802). All the data supporting the findings of this study are available from MetaboLights database or from the corresponding authors on reasonable request.

## Acknowledgments

This work was supported by the JDRF grants 4-1998-274, 4-1999-731 4-2001-435 and special research funds for Oulu, Tampere and Turku University Hospitals in Finland. This work was supported by the Juvenile Diabetes Research Foundation (2-SRA-2014-159-Q-R to M.O.) and the Academy of Finland (Centre of Excellence in Molecular Systems Immunology and Physiology Research - SyMMyS, Decision No. 250114, to M.O. and M.K.). We thank Olli Simell for his contribution in the DIPP study, to Anette Untermann for excellent technical support in metabolomics analysis, and to Alex Dickens and Partho Sen for helpful discussions and insight in relation to this study.

## Author contributions

M.O. and M.K. designed and supervised the study. K.T. and T.H. performed metabolomic analysis. S.L. and E.K. analysed the data. H.S., H.H., J.I., J.T. and R.V. contributed to the design and conduct of the clinical study. S.L. and M.O. wrote the manuscript. All authors critically reviewed and approved the final manuscript.

## Ethical approval and informed consent

The ethics and research committee of the participating university and hospital at University of Tampere, Tampere Finland, approved the study protocol. The study was conducted according to the guidelines in the Declaration of Helsinki. Written informed consent was signed by the parents at the beginning of the study for participation of their children enrolled in the study.

